# Molecular mechanism of a single stranded DNA-stimulated bacterial immune peptidase

**DOI:** 10.64898/2026.05.16.725639

**Authors:** Adarsh Calloji, Lydia R. Chambers, Kevin D. Corbett

## Abstract

Bacteria encode hundreds of immune pathways that protect host cells against infection by bacteriophages. While immune pathways possess exquisite mechanisms for self-regulation to avoid aberrant activation, many are also tightly regulated at the level of transcription. Many immune operons are regulated by CapP+CapH, a two-protein transcriptional regulator system that triggers immune operon expression in response to DNA damage, by sensing the presence of single-stranded DNA byproducts of DNA damage repair. Here we define how the CapP peptidase is activated by single-stranded DNA. DNA binding in a conserved inter-domain groove in CapP triggers rearrangement of the autoregulatory “cysteine switch loop”, opening the active site and allowing binding and cleavage of CapH, which in turn leads to transcriptional activation of an associated immune operon. Our data define a conserved molecular mechanism for sensing bacteriophage infection via DNA damage, and for triggering increased expression of immune operons in response.

## INTRODUCTION

Over billions of years, bacteria have evolved a diverse array of immune pathways that protect against infection by bacteriophages (phages), including CRISPR-Cas, restriction-modification, and cyclic oligonucleotide-based antiphage signaling systems (CBASS), among others^1–3^. Anti-phage immune pathways generally fall into two classes: those that directly recognize and destroy foreign DNA through their sequence (e.g. CRISPR-Cas)^4^, modifications (e.g. restriction-modification pathways)^5,6^, end structure (e.g. Shedu)^7,8^, or intracellular localization (e.g. SNIPE)^9^; and those that recognize later phage replication intermediates and either slow key metabolic pathways or kill the host cell outright to prevent phage replication. This second class of pathways is often termed “abortive infection” pathways^10–13^, and these pathways must be tightly regulated to prevent aberrant activation in the absence of infection.

While the majority of identified anti-phage immune operons are thought to be constitutively expressed in their host cells^14^, a subset are regulated by global signaling pathways, including quorum sensing in *Serratia, Pseudomonas*, and *Aliivibrio* ^15–17^, and the Gac/Rsm translational regulation system in Pseudomonads^18^. Recently, several families of transcriptional regulators have been discovered that are encoded adjacent to CBASS, BREX, and other immune operons, and regulate the expression of these operons in response to phage infection and/or DNA damage. These include the WYL domain transcription factor CapW/BrxR^19–22^, the kinase-substrate protein pair CapK+CapS^23^, and the peptidase-substrate protein pair CapP+CapH(Lau et al. 2022). In all three cases, these regulators recognize DNA damage by sensing single-stranded DNA (ssDNA) that arises as a byproduct of DNA repair. CapW is a transcriptional repressor, and ssDNA binding to its C-terminal WYL domain triggers a conformational change to release its N-terminal winged helix domain from DNA^19^. In CapK+CapS, ssDNA binding by CapK activates the enzyme to phosphorylate the HTH (helix-turn-helix) transcriptional repressor CapS, releasing it from DNA^23^. In CapP+CapH, ssDNA binding by CapP activates it to cleave the HTH transcriptional repressor CapH, releasing it from DNA^24^. For both CapK and CapP, how ssDNA binding activates the protein’s enzymatic activity is not known.

CapP and its substrate CapH are related to a pair of proteins that regulate expression of DNA damage response genes in *Deinococcus* species: the peptidase IrrE (also called PprI) and the HTH transcriptional repressor DdrO^25–29^. Similarly to CapH, DdrO forms a dimer and binds the promoters of certain DNA damage response genes, repressing their transcription^28^. IrrE cleaves DdrO in response to DNA damage, thereby releasing it from DNA to relieve transcriptional repression^30^. Recently, a structure of IrrE bound to ssDNA was determined by X-ray crystallography, revealing that ssDNA binds in a groove between the protein’s N-terminal zinc metallopeptidase domain and its C-terminal GAF domain, and packing against its central HTH domain^31^. ssDNA binding was found to promote DdrO binding and cleavage by IrrE^31^. CapP+CapH and IrrE+DdrO are also related to ImmA+ImmR systems, which control mobilization of prophages and ICE elements in response to DNA damage^32–34^.

Here, we combine protein structure prediction and biochemical assays to reveal the molecular mechanism of CapP activation by ssDNA. We show that CapP binds ssDNA equivalently to *Deinococcus* IrrE, in a groove between its N-terminal peptidase domain and its C-terminal GAF domain. In contrast to IrrE, our results suggest that ssDNA binding triggers rearrangement of CapP’s “cysteine switch loop,” disengaging this loop from the peptidase active site to activate the enzyme. Our modeling also reveals how CapP recognizes CapH through a hydrophobic surface on CapP that mimics the CapH dimerization interface, and positions CapH near the CapP active site for cleavage. Together, these findings provide a structural framework for understanding how CapP functions as a sensor that directly links DNA damage to transcriptional control.

## RESULTS

### CapP binds single-stranded DNA in a conserved interdomain groove

Our prior work showed that CapP binds single-stranded DNA (ssDNA), and that ssDNA binding activates CapP’s peptidase domain to cleave CapH^24^. To determine how CapP binds ssDNA, we first attempted to crystallize CapP from both *E. coli* MS115-1 and *Thauera* sp. K11 (56% identical to *E. coli* MS115-1) in the presence of ssDNA. Based on our previous observations showing that CapP preferentially binds poly-T DNA^24^, we attempted co-crystallization with a short poly-T ssDNA oligonucleotide, without success. As an alternative strategy, we performed AlphaFold 3 structure predictions with *E. coli* M115-1 CapP, both alone (with a bound Zn^2+^ ion; **Figure 1a-c** and **Figure S1**) and bound to a 12mer poly-T ssDNA and CapH (**Figure 1d-e** and **Figure S2**).

**Figure 1.**
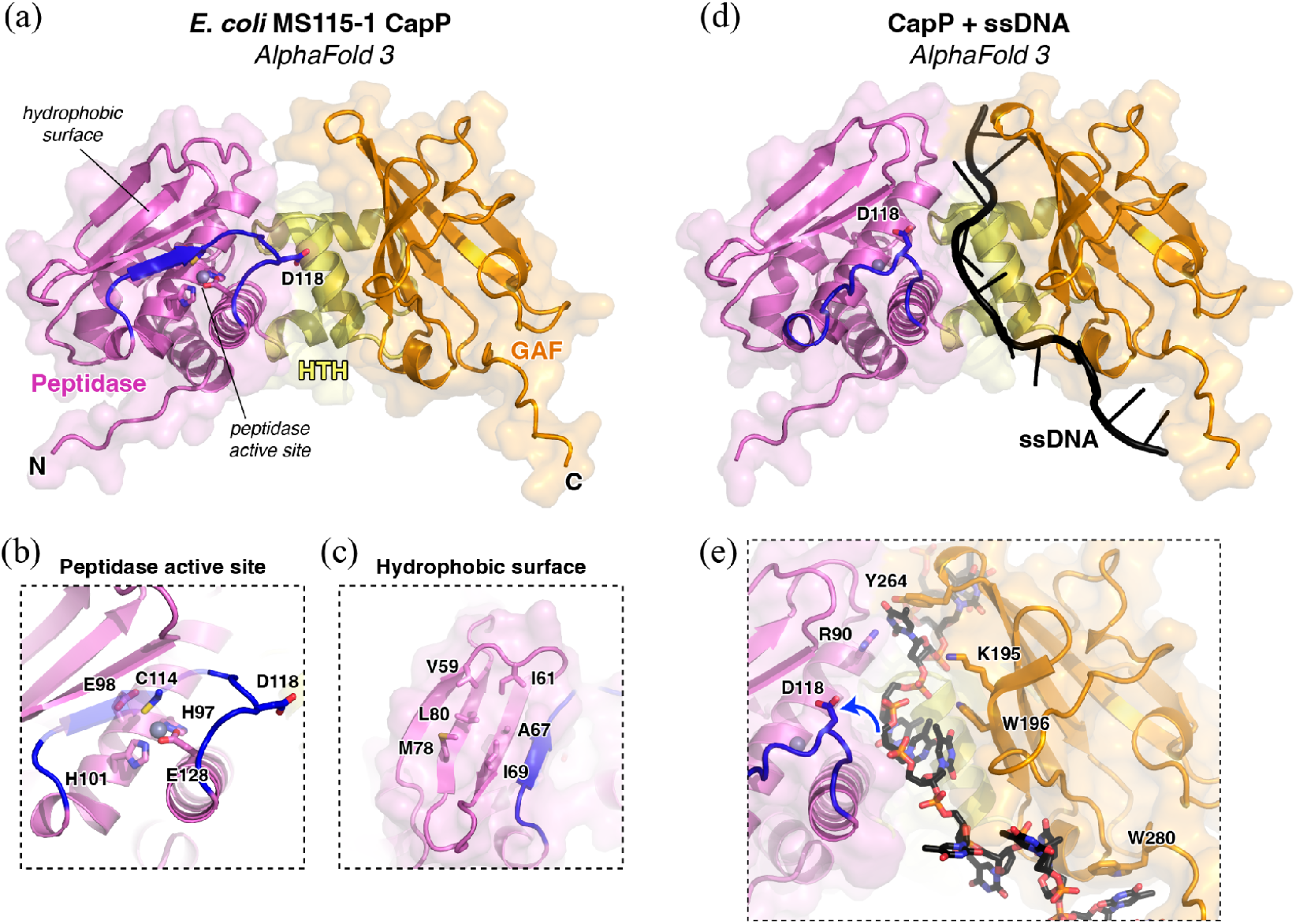
Predicted structures of *E. coli* CapP alone and bound to ssDNA. (a) AlphaFold 3 predicted structure of *E. coli* MS115-1 CapP bound to zinc (gray sphere). Domains are colored as follows: Peptidase (residues 1-141) magenta, HTH (residues 142-182) yellow, and GAF (residues 182-288) orange. The cysteine switch loop (residues 109-122) is colored blue). See **Figure S1** for details of structure prediction. (b) Closeup of the peptidase active site of *E. coli* CapP, showing conserved active site residues and residue D118 on the cysteine switch loop. (c) Closeup of the hydrophobic surface on the upper face of the CapP peptidase domain. (d) AlphaFold 3 predicted structure of CapP bound to ssDNA (12mer poly-T; black), from a prediction of CapP + CapH + ssDNA (see **Figure S2** for details). See **Figure S3** for comparison with *Deinococcus* IrrE bound to ssDNA. (e) Closeup of the predicted DNA binding groove of CapP, with residues mutated for biochemical assays shown as sticks and labeled. The prediction suggests that when CapP is bound to DNA, CapP’s cysteine switch loop swings outward to expose the active site (blue arrow).

The AlphaFold-predicted structure of *E. coli* MS115-1 CapP is extremely similar to our prior crystal structure of *Thauera* sp. K11 CapP (2.9 Å Cα r.m.s.d. over 272 residues)^24^. As in our prior structure, the CapP active site comprises three Zn^2+^-coordinating residues (H97, H101, and E128) plus a nearby catalytic glutamate residue (E98). The CapP peptidase domain contains an internal loop termed the “cysteine switch loop” with a conserved cysteine (C114 in *E. coli* MS115-1 CapP) that completes the coordination of the bound Zn^2+^ ion and is presumed to keep the enzyme inactive in the absence of ssDNA (**Figure 1b**). Not all CapP homologs possess the cysteine switch, but among those that do, the cysteine switch loop also contains a pair of well-conserved acidic residues (in *E. coli* MS115-1 CapP, D117 and D118) that are positioned close to the inter-domain groove between the protein’s N-terminal peptidase and C-terminal GAF domains (**Figure 1a,b, Figure S3**). Since this groove is where *Deinococcus* IrrE binds ssDNA^31^, we hypothesized that these acidic residues may be involved in sensing ssDNA; this idea is discussed further below. Finally, we observed that the CapP peptidase domain has a prominent hydrophobic surface adjacent to the active site, that in *E. coli* MS115-1 CapP comprises six solvent-exposed hydrophobic residues: V59, I61, A67, I69, M78, and L80 (**Figure 1c**).

Our AlphaFold-predicted structure of *E. coli* MS115-1 CapP bound to a poly-T ssDNA and CapH shows shows the ssDNA binding in a prominent groove in CapP between the protein’s N-terminal peptidase and C-terminal GAF domains, packing against the central HTH domain (**Figure 1d-e**). The predicted structure and binding pose of ssDNA is very similar to the crystal structure of *Deinococcus* IrrE bound to ssDNA (4.7 Å Cα r.m.s.d. over 160 residues; **Figure S4**)^31^. While the overall structure of CapP is largely identical in the AlphaFold predictions in the presence and absence of ssDNA, one major change is that in the presence of ssDNA, the cysteine switch loop is predicted to swing away from the bound ssDNA, positioning C114 more than 10 Å away from the active site Zn^2+^ ion (**Figure 1e**). In the predicted structure, several well-conserved positively-charged or aromatic residues in both the CapP peptidase and GAF domains are positioned close to the bound ssDNA, including R90, K195, W196, Y264, and W280. Several of these residues are structurally equivalent to DNA-binding residues in IrrE, including CapP R90 (IrrE R85), K195 (IrrE R207), and Y264 (IrrE R267) (**Figure S4b**).

To validate the predicted structure of CapP bound to ssDNA, we purified wild-type *E. coli* MS115-1 CapP (**Figure S5a**) and used fluorescence polarization to measure the binding of CapP to a short poly-T ssDNA oligonucleotide, finding a dissociation constant (*K*_*d*_) of 0.65 µM (**Figure 2, Figure S5b-c**). We next individually mutated all five predicted DNA-binding residues to alanine (R90A, K195A, W196A, Y264A, and W280A), and tested ssDNA binding. Four of the five point-mutants reduced ssDNA binding affinity (*K*_*d*_ = 1.2-8.7 µM), with only the K195A mutation having little effect on binding (*K*_*d*_ = 0.28 µM; **Figure 2b, Figure S5b-c**). Because no single point-mutant fully eliminated detectable DNA binding, we designed a double-mutant combining R90A and W196A. This double-mutant showed no detectable ssDNA binding in our fluorescence polarization assay (**Figure 2a-b**). Overall, these data support the AlphaFold prediction in which ssDNA binds CapP in the groove between the protein’s peptidase and GAF domains.

**Figure 2.**
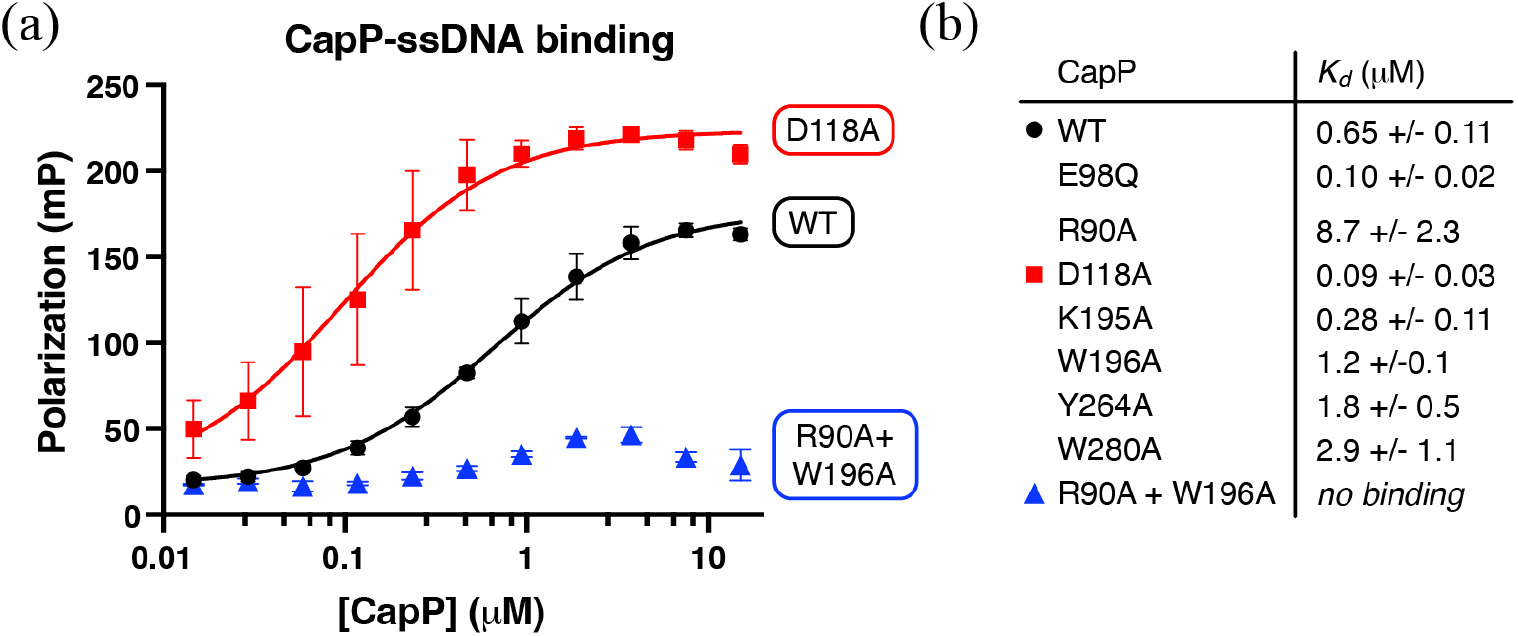
ssDNA binding by CapP. (a) Fluorescence polarization DNA binding assay for CapP wild-type (WT; black), D118A (red), and R90A+W196A mutant (blue) binding a 22mer poly-T ssDNA. (b) Table showing dissociation constants (*Kd*) for CapP WT and point mutants from fluorescence polarization assays (see **Figure S4** for protein purification, and **Figure S5** for binding curves).

We next mutated one of the conserved acidic residues on the CapP cysteine switch loop, D118, to alanine and tested ssDNA binding. We observed that the D118A mutant showed markedly tighter binding to ssDNA than wild-type CapP (*K*_*d*_ = 0.09 µM; **Figure 2b**). We hypothesize that this tighter ssDNA binding may arise from a loss of charge-repulsion effects when removing an acidic residue from the proximity of the ssDNA binding site.

### Molecular basis for CapH binding and cleavage by CapP

Our AlphaFold prediction of the CapP-ssDNA-CapH complex showed high confidence for CapP binding ssDNA (inter-chain ipTM (interface predicted template modeling score) = 0.80) and for CapP binding CapH (inter-chain ipTM = 0.82) (**Figure 3a, Figure S2**). In the prediction, CapH binds across CapP with its N-terminal DNA-binding HTH domain packing on the CapP GAF domain, and the C-terminal dimerization domain packing against the CapP peptidase domain. A highly-conserved tryptophan residue in the CapH HTH domain (W35) anchors the interaction with the CapP GAF domain, packing against CapP Y193 (**Figure 3c**). The predicted structure of the CapH C-terminal dimerization domain is significantly different from its native structure, which we previously determined by X-ray crystallography^24^. In the CapH homodimer, this domain forms two α-helices that dimerize in an interlocking “V” shape. In the predicted structure of a CapH protomer bound to CapP, the N-terminal α-helix of the CapH dimerization domain is predicted to unwrap, exposing the cleavage site (between F82 and R83)^24^ and positioning it near the active site (**Figure 3b**). The C-terminal α-helix maintains its helical structure and packs against the hydrophobic surface on the CapP peptidase domain, burying several hydrophobic residues that are also buried in the CapH homodimer (**Figure 3c-d**). Overall, this predicted structure points to a highly specific recognition of CapH by CapP, involving conserved residues on both CapH’s N-terminal HTH domain and its C-terminal dimerization domain. The interaction may also depend on ssDNA binding to CapP, which is predicted to open the cysteine switch loop and expose the peptidase active site for CapH binding. We attempted to directly detect CapP-CapH binding in the presence and absence of ssDNA using an affinity pulldown, without success (not shown). This result is consistent with reports that the binding of *Deinococcus* IrrE to its substrate DdrO is transient, even in the presence of ssDNA^31,35^.

**Figure 3.**
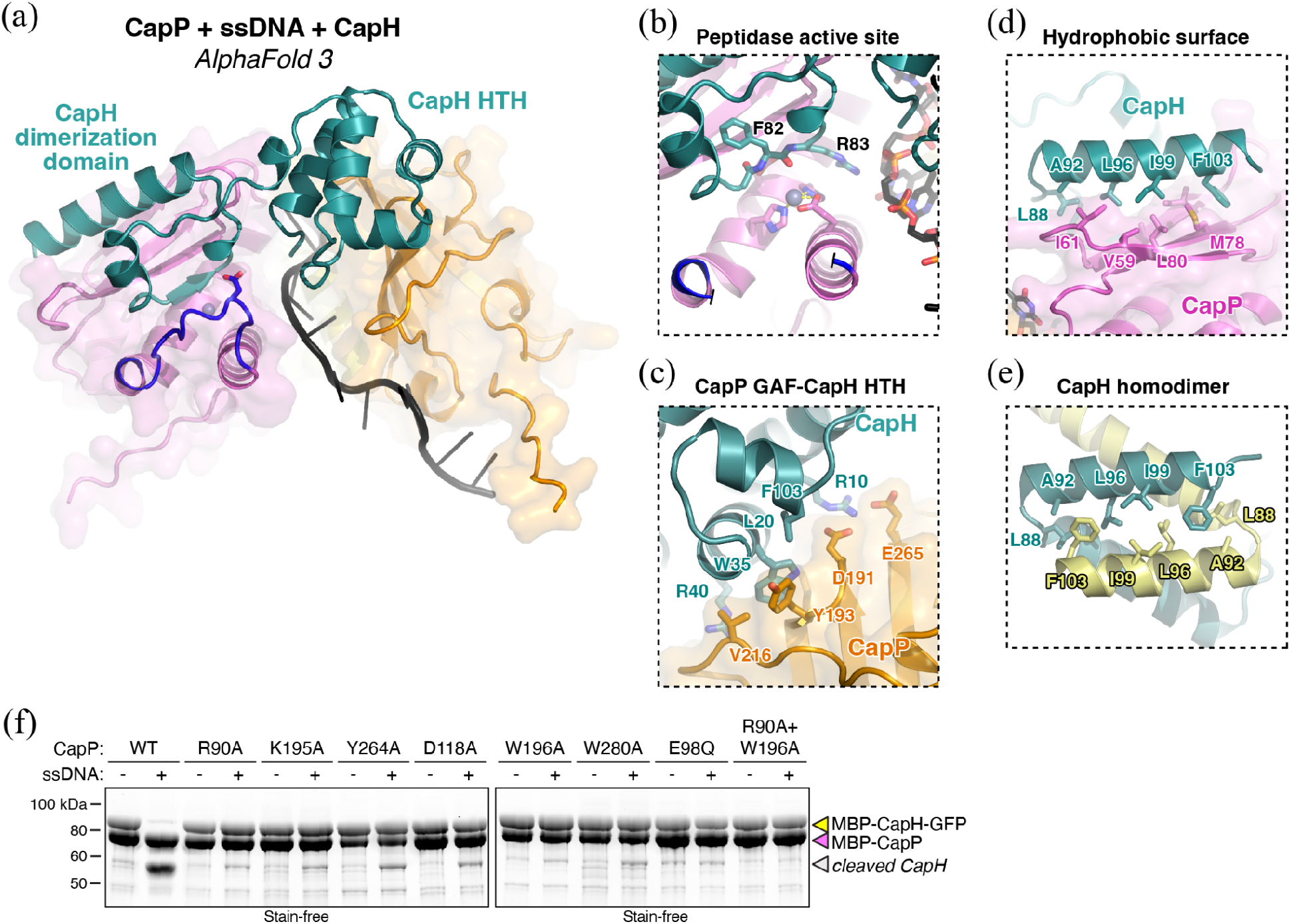
CapH cleavage by CapP. (a) AlphaFold 3 predicted structure of *E. coli* CapP + CapH + ssDNA. CapP is colored as in **Figure 1**, CapH is shown in dark green, and ssDNA is shown in black. See Figure S2 for structure-prediction details. (b) Closeup of the CapP active site in the predicted structure of *E. coli* CapP + CapH + ssDNA. The CapH cleavage site (F82-R83) is superimposed just above the bound zinc (modeled from the CapP+zinc predicted structure). The cysteine switch loop has been removed for visual clarity. (c) Closeup of the predicted interface between the CapP GAF domain and the CapH HTH domain, with interacting residues on each protein shown as sticks and labeled. (d) Closeup of the hydrophobic surface on the CapP peptidase domain, bound to the C-terminal α-helix of CapH. Interacting hydrophobic residues on both proteins are shown as sticks and labeled. CapP A67 and I69 (at rear) are not labeled. (e) View of the *E. coli* CapH C-terminal domain homodimer (PDB ID 7T5W)^24^, with the C-terminal α-helix of one protomer (dark green) oriented as in panel (c). (f) ssDNA-stimulated CapH cleavage by CapP wild-type (WT) and indicated point mutants. Activity is measured by the intensity of the cleaved CapH band (gray triangle). See **Figure S4** for SDS-PAGE analysis of individual proteins.

To test the functional linkage between ssDNA binding and CapH cleavage, we performed CapH cleavage assays with CapP point-mutants in the presence and absence of ssDNA. We previously used this assay to demonstrate that CapP cleaves CapH only in the presence of ssDNA, and that the active-site mutant E98Q shows no cleavage^24^. In this assay, we found that all five single point mutants we designed to compromise ssDNA binding (R90A, K195A, W196A, Y264A, and W280A) showed markedly reduced, but still detectable, CapH cleavage (**Figure 3e**). The reduced CapH cleavage in the CapP K195A mutant was surprising, given that this mutant did not affect ssDNA binding by CapP. Further inspection of the predicted CapP-CapH complex reveals that this residue comprises part of the predicted interface between the CapP GAF domain and the CapH HTH domain (**Figure S2d**); the K195A mutant may therefore affect CapH binding directly, rather than ssDNA binding. The CapP R90A+W196A double-mutant, which did not detectably bind ssDNA, also showed no detectable CapH cleavage (**Figure 3e**).

Strikingly, the CapP D118A switch loop mutant, which showed tighter ssDNA binding than wild-type CapP, also showed markedly reduced CapH cleavage activity compared to wild-type (**Figure 3e**). This result is consistent with a model in which D118 functionally couples ssDNA binding to movement of the cysteine switch loop; without this coupling, the active site remains occluded by the cysteine switch, preventing CapH cleavage.

## DISCUSSION

Here, we define the molecular mechanism of ssDNA sensing and peptidase activation in the CapP+CapH transcriptional regulator system, which is encoded adjacent to and regulates a range of bacterial immune operons^24^. We find that ssDNA binds in a conserved interdomain groove in CapP, mediating the disengagement of the protein’s cysteine switch from the active site to activate the CapP peptidase (**Figure 4**). We identify a set of key ssDNA-binding residues in the CapP peptidase and GAF domains, and identify acidic residues on the cysteine switch loop as essential for coupling ssDNA binding to peptidase activation. We also identify a prominent hydrophobic surface on the peptidase domain of CapP, which is predicted to bind the hydrophobic dimer interface in CapH to stabilize this protein in the monomer form and position it for cleavage by CapP.

**Figure 4.**
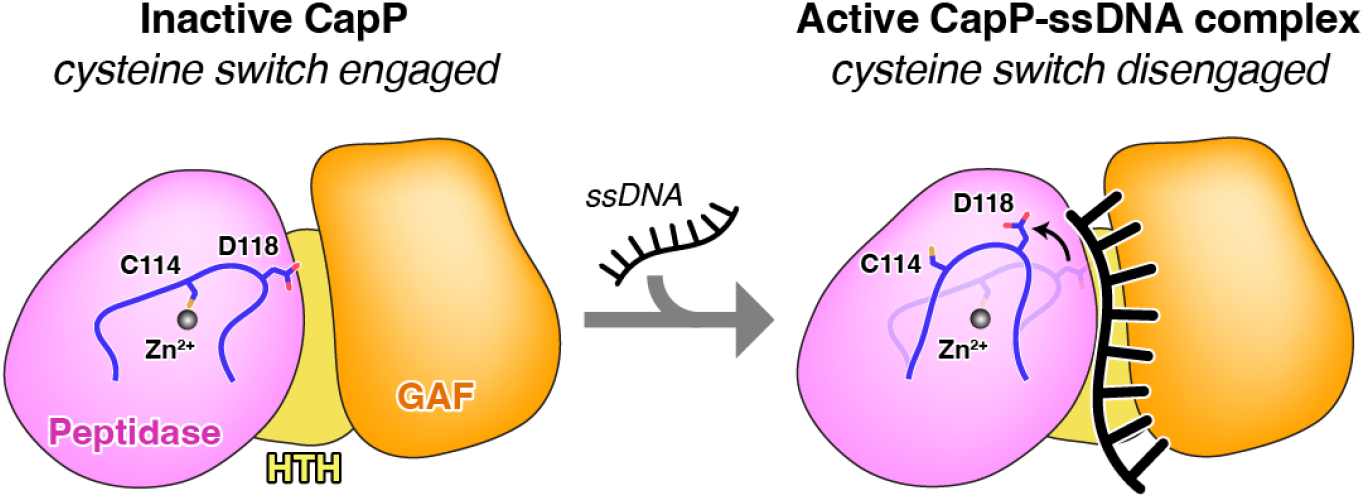
Model for CapP activation. Model for CapP activation by ssDNA, with D118 sensing binding of ssDNA to disengage the cysteine switch (blue) from the Zn^2+^ ion in the peptidase active site (black sphere).

Our findings regarding the ssDNA binding and activation mechanism of CapP closely mirror prior data on activation of the related peptidase IrrE, which participates in the DNA damage response in *Deinococcus. Deinococcus* IrrE shares a common domain structure with CapP^25^, likely binds its substrate DdrO through a similar peptidase-domain hydrophobic surface as we find in CapP (**Figure S4c** and ref. ^35^), and binds ssDNA in a groove between its peptidase and GAF domains^31^. IrrE lacks the cysteine switch of CapP, suggesting that its peptidase activity may be less tightly controlled than that of CapP. Indeed, prior biochemical studies have shown that purified IrrE cleaves DdrO *in vitro* without added ssDNA, but is further stimulated in the presence of ssDNA ^30,31^. In contrast, we previously showed that CapP is completely inactive in the absence of ssDNA^24^. In the absence of the cysteine switch, ssDNA-mediated stimulation of IrrE may occur simply via stabilization of DdrO binding in the presence of ssDNA^31,35^. In cells, IrrE activity has also been shown to depend on the level of free zinc ions in solution, which is potentially linked to exposure of cells to radiation and/or redox stress^36^.

A key remaining question in the field of bacterial immunity is why some immune pathways are regulated by DNA damage-responsive sensors, and some are not. For example, less than 10% of CBASS operons are associated with either CapP+CapH^24^ or CapW^20^; the rest are likely constitutively expressed. We have previously tested the ability of both CapW and CapP+CapH-associated CBASS operons to protect against lytic bacteriophage infection, and in both cases found more effective protection (as measured by plaque assays) when the operon is constitutively expressed, versus controlled by its native regulator(s)^20,24^. While we speculate that tight transcriptional regulation might provide a long-term survival benefit by preventing aberrant activation and cell death in the absence of lytic phage infection, this model has not been rigorously tested. It is also possible that these immune pathways specifically combat integration or excision of lysogenic phages, by piggybacking on the stress signals that regulate these phages’ life cycle decisions.

Across the highly diverse immune repertoire of bacteria, proteases are emerging as key components that play a range of roles including infection sensing, activation of effector proteins, and serving as effectors themselves. We have shown that CapP recognizes ssDNA produced as a result of phage infection and, through its cleavage of CapH, triggers increased expression of diverse immune operons ^24^. *E. coli* Lit (Late inhibitors of T4) serves as both an infection sensor and effector, recognizing a complex of phage-encoded major capsid protein and the host Ef-Tu elongation factor, then cleaving Ef-Tu to kill the host and prevent phage propagation^37–40^. Recent comprehensive work from Gao and colleagues has identified a large set of protease infection sensors that cleave diverse operonic effector proteins, converting them from inactive zymogen forms to active effectors^41^. Proteases also function as effectors, for example caspase-like effectors in Type IV Thoeris^42^ and predicted protease domains in some Retron and Lamassu systems^43,44^. Given the tendency for novel immune pathways to evolve in a modular fashion, mixing and matching sensor and effector proteins and domains, we expect that more such systems – potentially involving CapP-related proteins activated by ssDNA – to be identified in the future.

## MATERIALS AND METHODS

### Protein structure predictions

All protein structure predictions were performed using AlphaFold 3 (ref. ^45^) using the AlphaFold Server (http://alphafoldserver.com). Structure comparisons and figure generation were performed using PyMOL version 3 (Schrödinger).

### Cloning and Protein Purification

Expression vectors for *E. coli* MS115-1 CapH (NCBI ID WP_001515173.1) and *E. coli* MS115-1 CapP (NCBI ID WP_001290439.1) were cloned as described^24^. CapH was cloned with an N-terminal His_6_-maltose binding protein (MBP) tag and a C-terminal green fluorescent protein (GFP) tag, and CapP was cloned with an N-terminal His_6_-MBP tag. CapP point mutants were generated with PCR-based mutagenesis.

For protein expression, plasmids were transformed into *E. coli* strain Rosetta 2 (DE3) pLysS (EMD Millipore) and grown in LB media with appropriate antibiotics. 5 mL of saturated overnight starter culture was used to inoculate 1 liter of 2XYT media and grown at 37°C to OD_600_=0.6, then induced with 0.25 mM Isopropyl-β-D-thiogalactopyranoside (IPTG) and grown a further 15 hours at 20°C. Cells were harvested by centrifugation and resuspended in Buffer A (20 mM Tris pH 7.5, 10% glycerol) plus 300 mM NaCl, 10 mM imidazole, and 5 mM β-mercaptoethanol (10 μM ZnCl_2_ was added to buffers for CapP purification). For purification, resuspended cells were lysed by sonication (Branson Sonifier), centrifuged, and clarified lysate was passed over a Ni^2+^ affinity column (Ni-NTA Superflow; Qiagen). Nickel columns were washed with buffer A plus 300 mM NaCl and 20 mM imidazole pH 8.0, then eluted in buffer A plus 300 mM NaCl and 400 mM imidazole. Protein was then passed over an anion-exchange column (Hitrap Q HP, Cytiva) in Buffer A with a 75 mM to 1 M NaCl gradient, and peak fractions were collected, concentrated, then passed over a size exclusion column Superdex 200, Cytiva) in buffer GF (buffer A plus 300 mM NaCl and 1 mM dithiothreitol (DTT)). Peak fractions were concentrated by ultrafiltration (Amicon Ultra, EMD Millipore) to 5-15 mg/mL and flash frozen in liquid nitrogen before being stored at -80°C.

### DNA Binding Assays

For ssDNA binding assays, triplicate 60 μL reactions containing MBP-CapP (wild type or indicated mutants), starting at 15 μM followed by two-fold dilutions, plus 50 nM 5′-FITC labeled 22mer poly-T DNA, in FP buffer (25 mM Tris pH 8.5, 50 mM sodium glutamate, 5 mM MgCl_2_, 1 mM DTT, 5% glycerol, 0.01% NP40 substitute) were mixed, then transferred to a 384-well microplate and incubated for 10 minutes at room temperature. Fluorescence polarization was read in a plate reader (Tecan Spark), and data were analyzed in GraphPad Prism using a single-site binding model.

### CapP cleavage assays

For CapH cleavage assays, 10 μl reactions containing 10 μM MBP-CapP (wild type or indicated mutants), 10 μM MBP-CapH-GFP, and 10 μM 22mer poly-T ssDNA in cleavage buffer (50 mM Tris pH 7.5, 50 mM NaCl and 5 μM ZnCl_2_) were mixed, then incubated at 37°C for 2 hours.

Samples were separated by SDS-PAGE and visualized using Stain-Free imaging (Bio-Rad).

## ACKNOWLEDGEMENTS

The authors thank members of the Corbett lab for mentorship and helpful conversations. The authors acknowledge support from the National Institutes of Health (R35 GM144121 to K.D.C., T32 GM139795 to L.R.C, and F31 AI194603 to L.R.C.).

## AUTHOR CONTRIBUTIONS

A.C. performed protein structure prediction, molecular cloning, protein purification, and biochemical assays, and wrote the paper. L.R.C. performed protein purification and biochemical assays. K.D.C. designed the project, oversaw experiments, generated figures, and wrote the paper.

## DECLARATION OF INTERESTS

The authors declare no competing interests.

## SUPPLEMENTAL FIGURES

**Figure S1.**
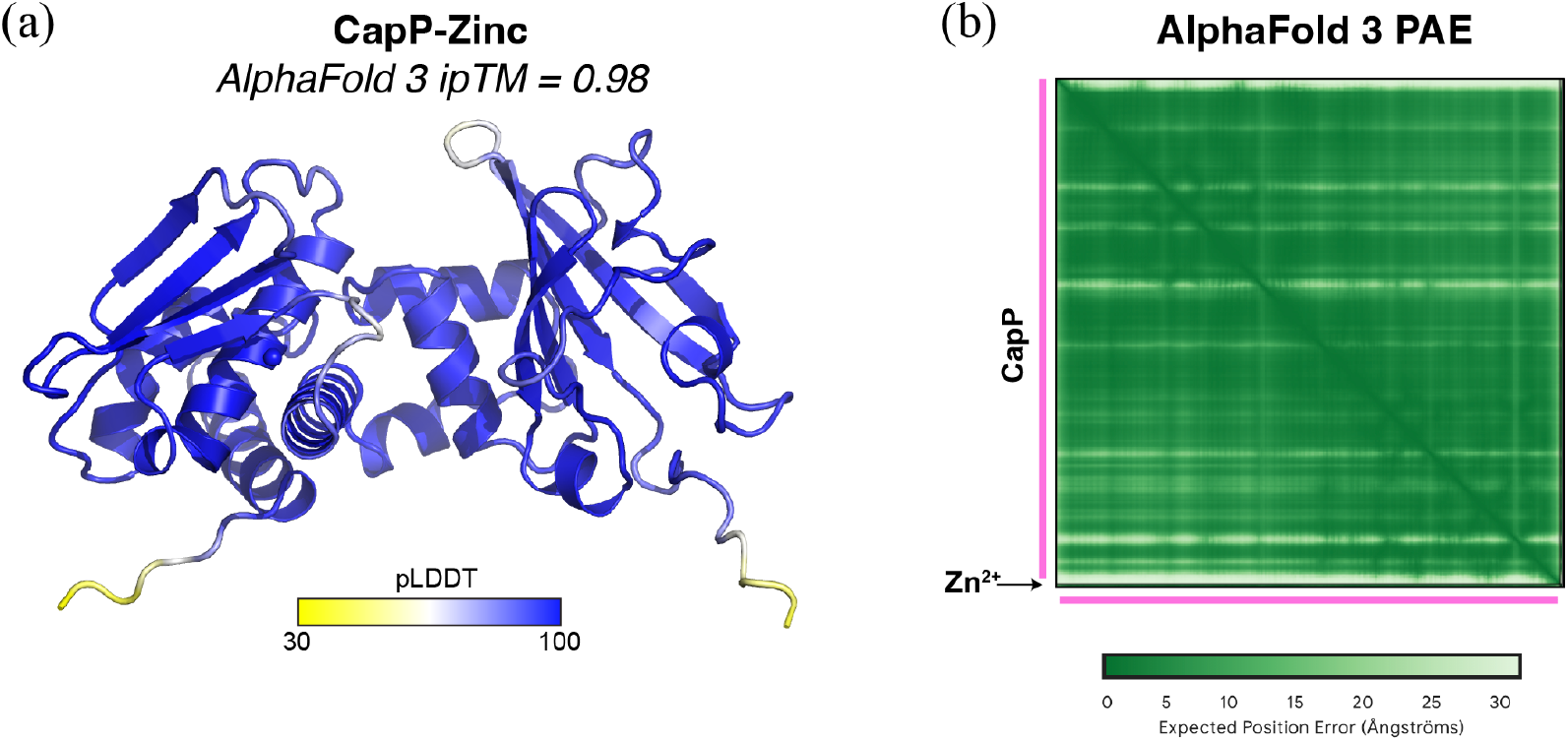
Predicted structure of *E. coli* CapP. (a) AlphaFold 3 predicted structure of *E. coli* MS115-1 CapP bound to a zinc ion, colored by local confidence (pLDDT). (b) AlphaFold 3 predicted aligned error (PAE) plot for the prediction shown in panel (a).

**Figure S2.**
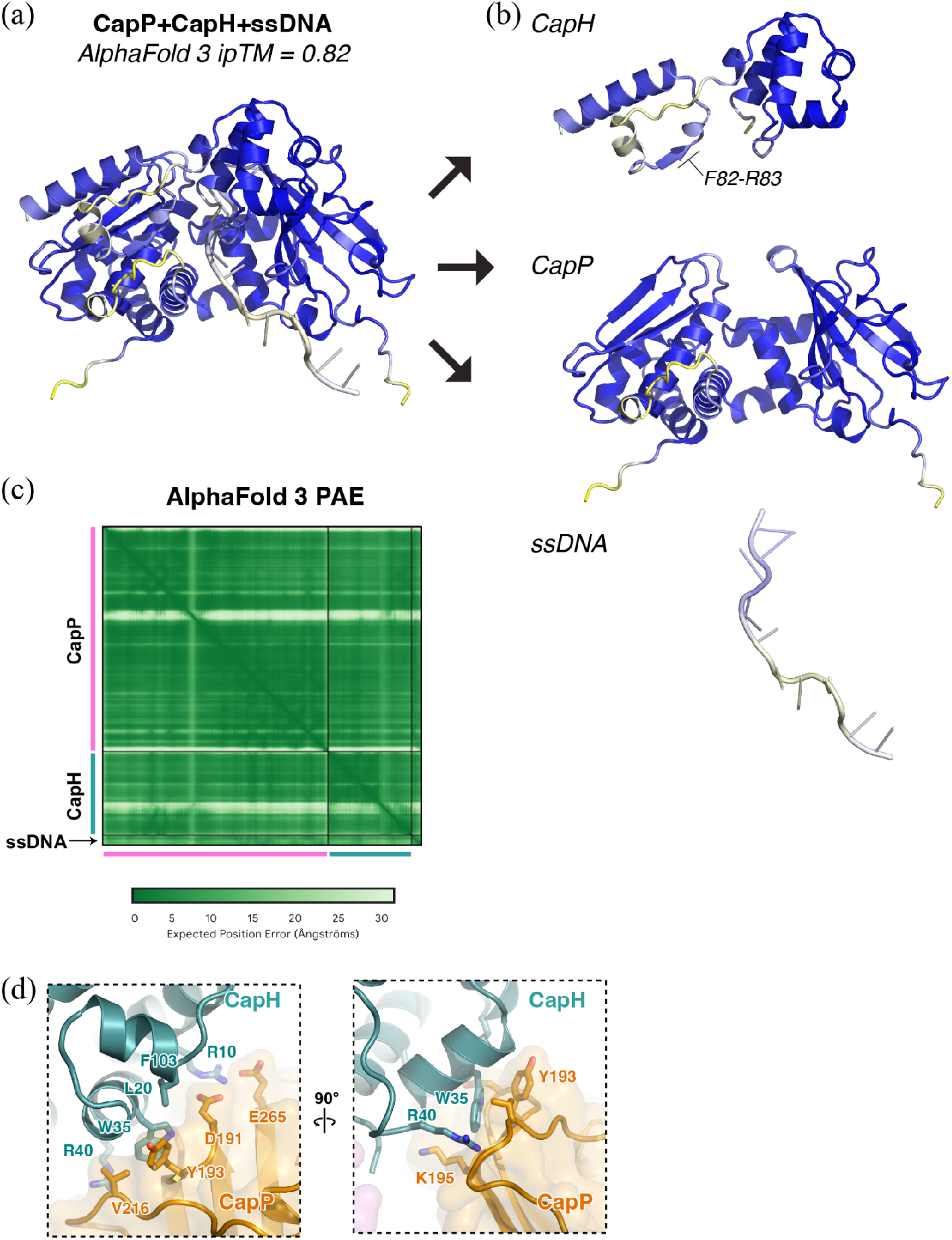
Predicted structure of *E. coli* CapP + CapH + ssDNA. (a) AlphaFold 3 predicted structure of *E. coli* MS115-1 CapP + CapH + ssDNA (12mer poly-T), colored by local confidence (pLDDT). (b) View as in panel (a), separated by macromolecular component: CapH top, CapP middle, and ssDNA bottom. For CapH, the cleavage site (F82-R83; labeled) is modeled with high confidence (low pLDDT). (c) AlphaFold 3 predicted aligned error (PAE) plot for the prediction shown in panel (a). (d) Two views of the predicted interaction between the CapH N-terminal HTH domain (green) and the CapP GAF domain (orange). The left view is equivalent to **Figure 3c**; the right view shows a rotated view with CapP K195 visible.

**Figure S3.**
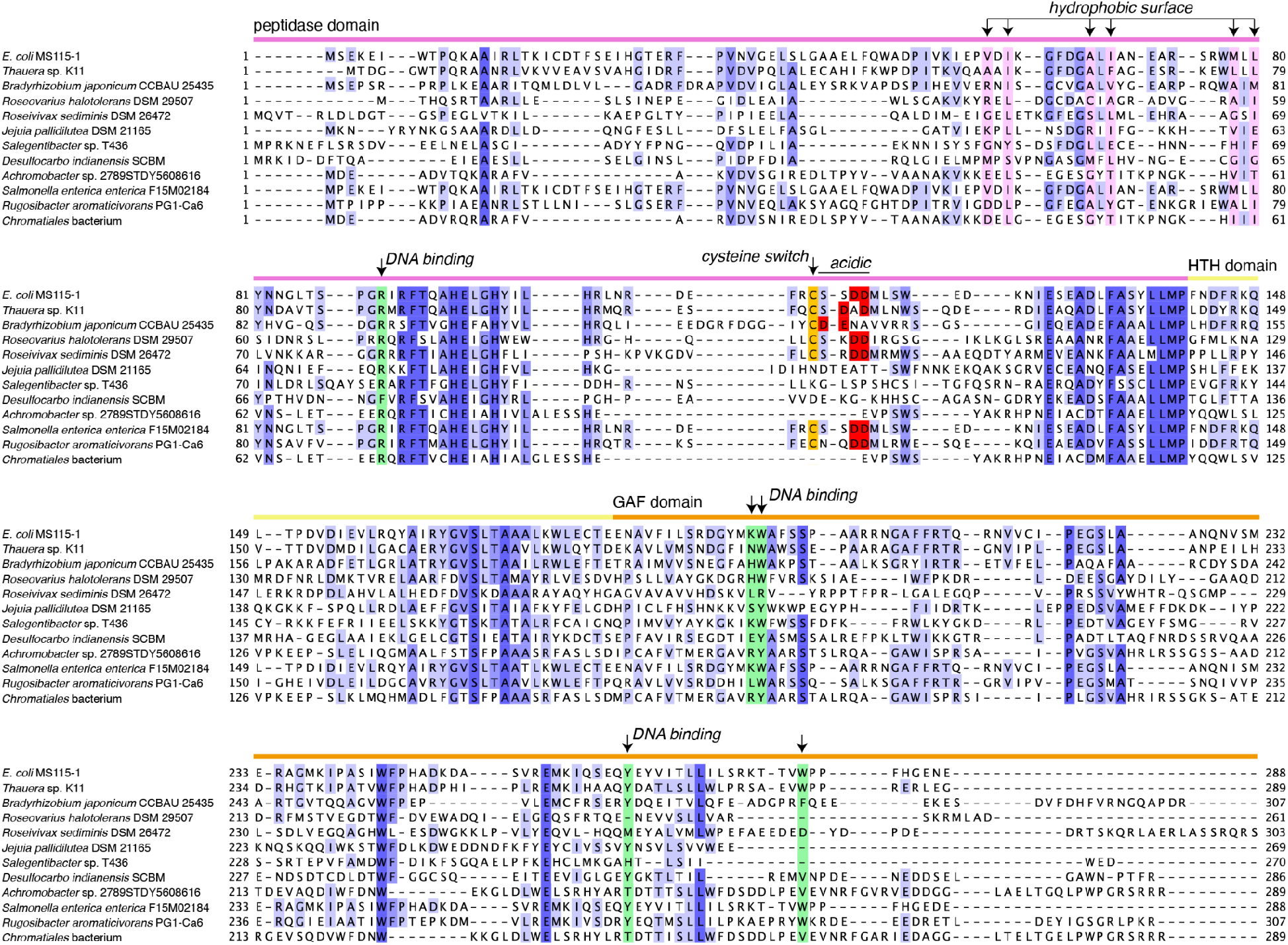
CapP sequence alignment. Protein sequence alignment for selected CapP homologs identified previously^24^. Domains are noted and colored as in **Figure 1**. Residues involved in the peptidase-domain hydrophobic surface are indicated and highlighted in pink; residues predicted to interact with ssDNA are indicated and highlighted in green. Cysteine switch residues are highlighted in orange, and acidic residues on the cysteine switch loop are highlighted in red. Accession numbers for each sequence are as follows: *E. coli* MS115-1 (NCBI WP_001290439.1), *Thauera* sp. K11 (Uniprot A0ACD6B902), *Bradyrhizobium japonicum* CCBAU 25435 (IMG 2702575792), *Roseovarius halotolerans* DSM 29507 (IMG 2730460223), *Roseivivax sediminis* DSM 26472 (IMG 2635168333), *Jejuia pallidilutea* DSM 21165 (IMG 2729910569), *Salegentibacter* sp. T436 (IMG 2719648675), *Desulfocarbo indianensis* SCBM (IMG 26548996400), *Achromobacter* sp. 2789STDY5608616 (IMG 2678852882), *Salmonella enterica enterica* F15M02184 (IMG 2744296958), *Rugosibacter aromaticivorans* PG1-Ca6 (IMG 2595015096), *Chromatiales* bacterium (IMG 2502333868).

**Figure S4.**
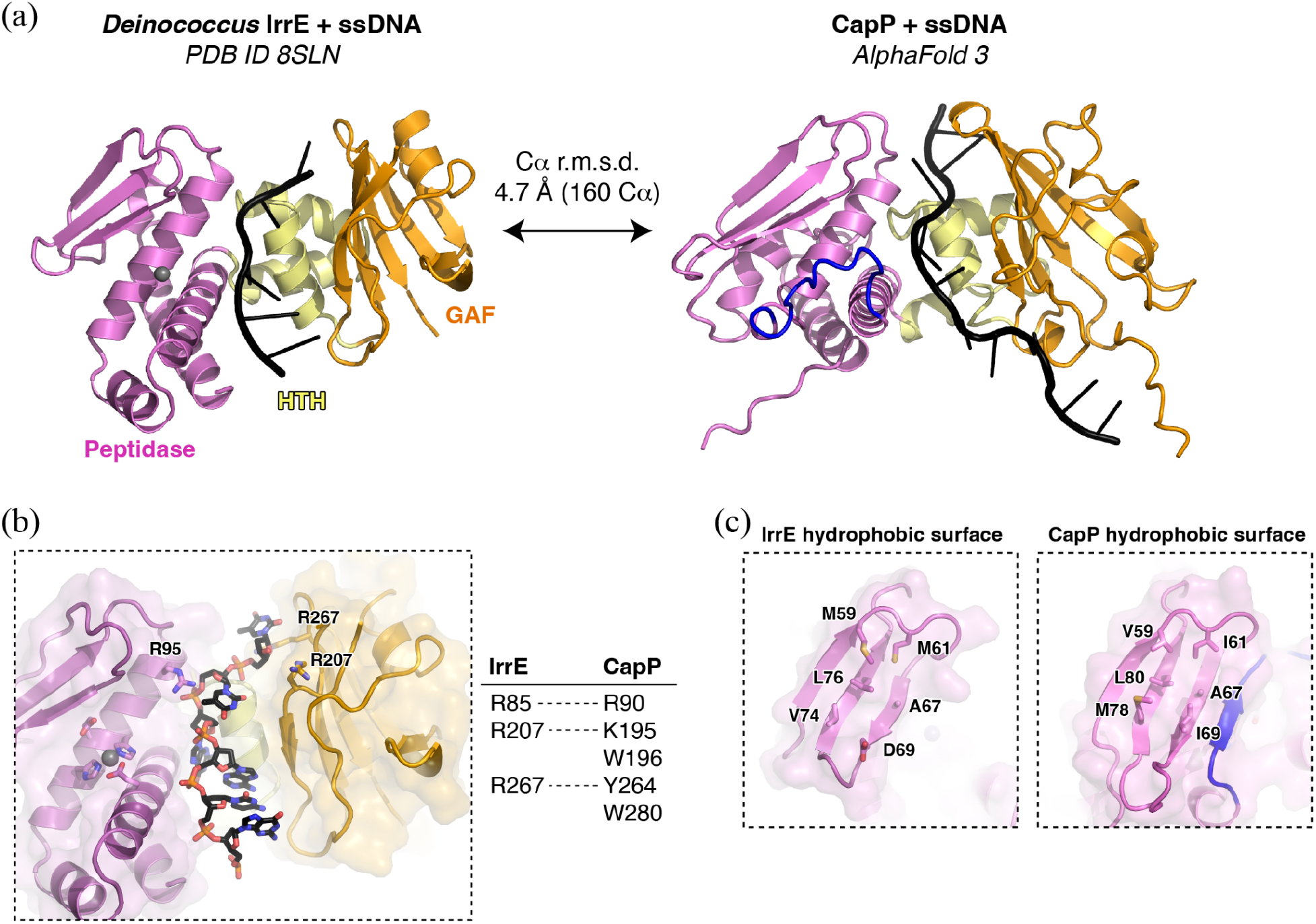
Comparison of ssDNA-bound IrrE and CapP. (a) *Left:* Crystal structure of *Deinococcus* IrrE bound to ssDNA (PDB ID 8SLN)^31^, with Peptidase, HTH, and GAF domains colored and labeled, and ssDNA shown in black. *Right:* Predicted structure of *E. coli* CapP bound to ssDNA (see **Figure S2**). (b) Closeup of *Deinococcus* IrrE ssDNA binding, with residues whose mutation was shown to affect ssDNA binding shown as sticks and labeled. *Right:* table showing candidate DNA-binding residues in IrrE and CapP that occupy structurally equivalent positions. (c) *Left:* Closeup of the hydrophobic surface on the *Deinococcus* IrrE peptidase domain. *Right:* Closeup of the hydrophobic surface on the *E. coli* CapP peptidase domain.

**Figure S5.**
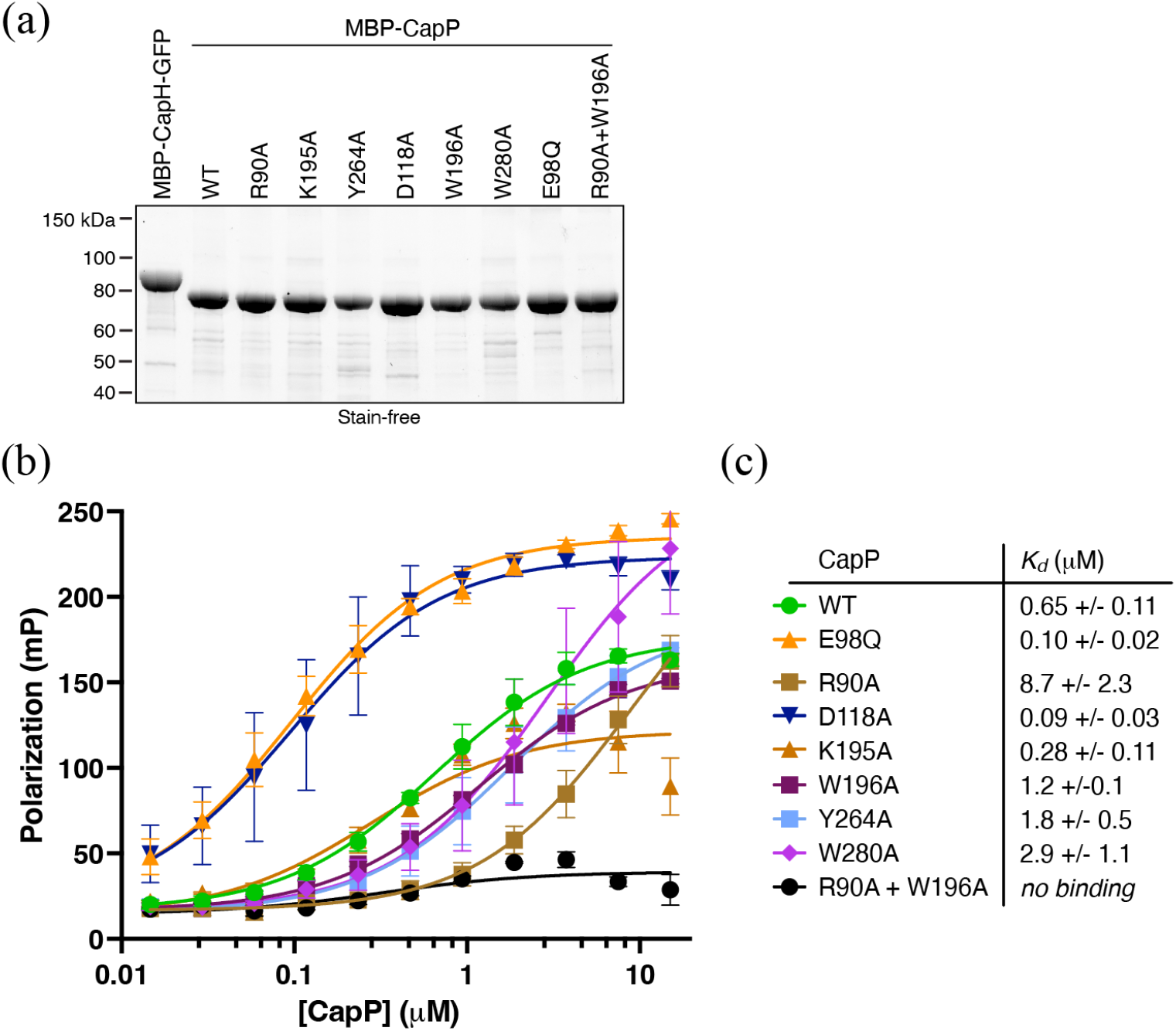
Purification and ssDNA binding by CapP and CapP point mutants. (a) SDS-PAGE analysis of all purified proteins used for biochemical assays. (b) Fluorescence polarization DNA binding assay for CapP wild-type (WT; green) and point mutants (key in panel (b)) binding a 22mer poly-T ssDNA. (c) Table showing dissociation constants (*Kd*) for CapP WT and point mutants from fluorescence polarization assays.

## Notes

### Competing Interest Statement

The authors have declared no competing interest.

